# Privacy-Preserving Read Mapping Using Locality Sensitive Hashing and Secure Kmer Voting

**DOI:** 10.1101/046920

**Authors:** Victoria Popic, Serafim Batzoglou

## Abstract

The recent explosion in the amount of available genome sequencing data imposes high computational demands on the tools designed to analyze it. Low-cost cloud computing has the potential to alleviate this burden. However, moving personal genome data analysis to the cloud raises serious privacy concerns. Read alignment is a critical and computationally intensive first step of most genomic data analysis pipelines. While significant effort has been dedicated to optimize the sensitivity and runtime efficiency of this step, few approaches have addressed outsourcing this computation securely to an untrusted party. The few secure solutions that have been proposed either do not scale to whole genome sequencing datasets or are not competitive with the state of the art in read mapping. In this paper, we present **BALAUR**, a privacy-preserving read mapping algorithm based on locality sensitive hashing and secure kmer voting. **BALAUR** securely outsources a significant portion of the computation to the public cloud by formulating the alignment task as a voting scheme between encrypted read and reference kmers. Our approach can easily handle typical genome-scale datasets and is highly competitive with non-cryptographic state-of-the-art read aligners in both accuracy and runtime performance on simulated and real read data. Moreover, our approach is significantly faster than state-of-the-art read aligners in long read mapping.

## 1 Introduction

Recent sequencing technology breakthroughs have resulted in a dramatic increase in the amount of available sequencing data, enabling important scientific advances in biology and medicine. At the same time, the compute and storage demands associated with processing genomic data have also substantially increased and now often outmatch the in-house compute capabilities of many research institutions. Outsourcing computation to commercial low-cost clouds (e.g., Amazon Elastic Compute Cloud, Azure, Google Cloud), which offer the ability to allocate massive compute power and storage on demand, provides a convenient and cost-effective solution to this problem. However, exposing genomic data to an untrusted third party also raises serious privacy concerns since genomic data carries extremely sensitive personal information about its owner, such as ethnicity, ancestry, and susceptibility to certain diseases. A recent review [9] describes the growing concern over the ability to protect personal genetic privacy. As summarized by the review, the privacy of genetic information is currently a demand of many regulatory statutes in the US and EU and a major determinant of whether individuals are willing to participate in scientific studies. While, de-identification (i.e., removal of personal identifiers) and anonymization techniques have been suggested as solutions to this problem, it has been shown that such techniques cannot reliably prevent the identification of an individual from genomic data [9,12,31,33]. For example, numerous identity-tracing attacks have been demonstrated using quasi-identifiers, including demographic metadata [29,30], pedigree structures [25], and genealogical data [11,17]. The vulnerabilities of such approaches motivate the need for cryptographic techniques to process and store genomic information.

Read alignment is a critical and computationally intensive first step of most genomic data analysis pipelines. While tremendous effort has been dedicated to this problem, leading to the development of many highly efficient read mapping tools [18,20,21,23], few approaches have addressed outsourcing this computation securely to an untrusted party. The few secure solutions that exist either do not scale to whole genome sequencing [3, 13, 15] datasets or are not competitive with the state of the art in read mapping [8]. For example, the protocol [3] for computing edit distances using homomorphic encryption (HOM) [10] requires 5 minutes on a single pair of 25-bp sequences [15], and the approach [13] using secure multi-party computations, while more efficient, still takes 4 seconds for a pair of 100-bp sequences. Such compute costs cannot scale to whole genome computations, which require the pairwise comparison of millions to billions of sequences. Recently, Chen et al [8] proposed a secure seed-and-extend read mapping algorithm on hybrid clouds, which splits the computation such that the public cloud finds the exact seed matches using encrypted seeds and the private cloud extends the seed matches using unencrypted data. With this approach, mapping 10 million 100-bp reads takes 372 CPU hours: 1.5h on 30 nodes (with 8 cores/node) on the public cloud and an additional 2h on the private cloud. Here the time spent on the private cloud alone is comparable with the total runtime of a standard state-of-the-art aligner. For instance, the Bowtie2 aligner [18] takes 12 minutes to map 2 million 100-bp long reads on a single core with a peak memory footprint of 3.24GB. Moreover, to align 100-bp long reads with an edit distance of six, this approach requires 6.8TB to store the reference index. Therefore, although it can handle genome-scale computations in reasonable time, this approach still significantly exceeds the compute and storage requirements of standard read aligners.

In this paper we introduce **BALAUR**, a novel privacy preserving read mapping technique for hybrid clouds that securely outsources a significant portion of the read-mapping task to the public cloud, while being highly competitive with existing state-of-the-art aligners in speed and accuracy. At a high level, **BALAUR** can be summarized in the following two phases: (1) fast identification of a few candidate alignment positions in the genome using the locality sensitive hashing (LSH) (on the secure private client) and (2) evaluation of each candidate using secure kmer voting (on the untrusted public server). We leverage LSH and the MinHash [7] technique in Phase 1, by formulating the task of finding the candidate alignment positions as nearest neighbor search under the Jaccard set similarity criterion, searching for reference sequences that are most similar to the reads. To rapidly find such sequences, we first pre-compute the MinHash fingerprints of reference genome windows and store them in our MinHash reference genome index (MHG) data structure, designed to efficiently scale to full human genome datasets. The high selectivity of the MHG is crucial for our alignment algorithm, since it significantly decreases the overhead associated with encrypting and transferring the selected reference sequences for secure voting in Phase 2. It also demonstrates the effectiveness of LSH and MinHash for whole-genome alignment in general and especially for long read mapping. We describe each step in detail in the remainder of this paper, analyze the security guarantees of our approach and present an evaluation of its performance on simulated and real read datasets. The **BALAUR** algorithm was implemented in *C*++ and is freely available at http:://viq854.github.com/balaur.

## 2 Background

### 2.1 Locality Sensitive Hashing

Locality sensitive hashing (LSH) is a probabilistic dimensionality-reduction technique that has been introduced by Indyk and Motwani [14] to address the approximate similarity search problem in high dimensions. Its key property is to maximize the probability of collision of objects that are similar. In particular, let *H* be a family of hash functions *h*: *R*^*d*^ ← *U*. An LSH scheme defines a probability distribution over the family *H*, such that given two objects *x,y* ∈ *R*^d^: *Pr*_*h∈H*_ [*h*(*x*) = *h*(*y*)] = *similarity*(*x,y*).

For example, a simple family of functions *H* can be constructed for d-dimensional binary vectors from {0, 1}^d^ under the Hamming distance metric. In this case, the family of functions can just consist of all the projections of the input points from {0, 1}^d^ onto one of the *d* vector coordinates; namely, of the functions *h*_*i*_(*x*) = *x*_*i*_, where *i* ∈ {1,…, *d}* is a random index into the vector *x*. It can be easily seen that under such hash functions, the probability of collision of the hashes of two given vectors will be equal to the fraction of coordinates that are equal between the two vectors. For a survey of different LSH families see [2]. In computational biology, LSH has been applied to several tasks, including motif discovery [32], genome-wide association studies [6], and more recently SMS read overlap detection for de novo assembly [5].

In this work we apply LSH to hash the reference genome and the reads, such that the hash values (referred to as *fingerprints*) of the read and its genome window collide with high probability. Since the read can differ from the reference sequence it maps to due to sequencing errors and true genomic variants, our similarity measure needs to handle differences in the two sequences arising from base substitutions and indels. A standard approach for measuring the similarity between two strings is to represent them as sets (e.g., a set of all the words in a document). Then the similarity of two such sets *A* and *B* can be expressed by their *Jaccard coefficient:* 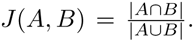. Several LSH families have been proposed for the Jaccard similarity criterion. Below we describe one of the most popular such techniques; namely, the MinHash algorithm, which we applied to NGS datasets in this work.

#### MinHash Algorithm

The *min-wise independent permutations* (MinHash) LSH family has been proposed by Broder et. at [7] for the Jaccard similarity measure and is defined as follows. Let *U* be the ground set of all possible set items. Given a random permutation π of indices of *U* and a set *X*, let *h*_π_(*X*) = *min*_*x∈X*_{π(*x*)}. The MinHash LSH family *H* will consist of all such functions for each choice of *π*. It can be easily shown that for a given *h* chosen uniformly at random, *Pr*[*h_π_*(*A*) = *h*_π_(*B*)] = *J* (*A, B*) (see [7] for details).

Due to the high variance in the above probability of collision, an amplification process is usually applied to reduce it. More specifically, instead of using one hash function, we concatenate *L* different hash functions from the family *H* chosen independently at random. It can be shown that given the number of the hash collisions among the chosen *L* functions, c, the ratio *c/L* can also be used as an unbiased estimator for *J* (*A, B*). Since computing random permutations can be prohibitive, the hash functions are typically created using universal hash functions of the form: *h*(*x*) = *ax* + *b*. We follow a similar approach in our method.

## 3 Methods

### 3.1 MinHash Fingerprinting

In order to apply the set Jaccard similarity measure to the reference genome windows and the read sequences, we represent them as sets of overlapping *kmers*. Given a sequence *S*, the set of overlapping kmers consists of all the substrings of *S* of length *k*. There is a clear tradeoff between the length *k* and the sensitivity of the results. If *k* is low, the resulting kmers will be short and shared by many sequences; on the other hand, if *k* is too large, there might be no kmer matches between the read and its corresponding reference window due to sequencing errors and genomic variations. We found k = 16 to provide a good balance. Furthermore, due to the repetitive nature of the genome, some kmers can be more informative than others. In particular, kmers that occur frequently across the genome carry little information value and are ineffective sequence markers. These kmers are analogous to uninformative commonplace words, such as ‘the’ and ‘an’, in a word document. We discard such frequently occurring kmers from the sequence set.

Given the set *K* = {*s*_0_, *s*_1_,…, *s*_n-1_} of kmers of length *k* of a read or reference window sequence *S*, its MinHash fingerprint vector *F* = [ƒ_0_,ƒ_1_,…,ƒ_L-1_] is created as follows. First each kmer in the set is hashed using a deterministic hash function *Ĥ*, resulting in the set of kmer hash values *K*_*Ĥ*_ = {*Ĥ*(*s*_0_), *Ĥ*(*s*_1_),…, *Ĥ*(*s*_n-1_)}. We then apply *L* random universal hash functions of the form *h*_*i*_(*x*) = *a*_*i*_*x* + *b*_*i*_ to each element of *K*_*Ĥ*_ The fingerprint entry ƒ_*i*_ is the minimum set element under each hash function *h*_g*i*_:

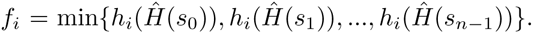

#### Algorithm 1 Rolling MinHash

1: procedure MinHashRoll(*M*, *m*_*oldest*_,*F*_*w*_)

2: *w*_*p+1;last*_ ← last kmer in window *w*_*p+1*_

3: *H*_*last*_(*w*_*p+1;last*_)

4: for *i* = 0 to *L* do

5: *min*_*h*_ ← *h*_*i*_(*H*_*last*_)

6: if *min*_*h*_ < *F*_*w*_*p*__ (*i*) then

7: *F*_*w*_(*i*) *min*_*h*_

8: *M*(*i,m*_*oldest*_) *min*_*h*_

9: else if *M*(*i*,_*moldest*_) != *F*_*w*_(*i*) then

10: *M*(*i*;*m*_*oldest*_) *min*_*h*_

11: else

12: *F*_*w*_(*i*) ← MAX_–_VAL

13: *M*(*i,m*_*oldest*_) ← *min*_*h*_

14: for *j* = 0 to *n* do

15: if *M*(*i; j*) < *F*_*w*_(*i*) then

16: *F*_*w*_(*i*) *M*(*i, j*)

17: *m*_*oldest*_ (*m*_*oldest*_ + 1) mod *n*

### 3.2 MinHash Reference Genome Index

To rapidly find the windows of the genome most similar to the reads under the Jaccard metric, we construct a MinHash reference genome index (MHG) as follows. We apply the MinHash algorithm to each window of the genome (of length equal to the length of the reads) to obtain its fingerprint *F*_*w*_. Windows of the genome containing only high-frequency kmers and ambiguous-base kmers are excluded. Once the fingerprints are computed, each window is stored in multiple tables corresponding to distinct fingerprint projections. We describe these steps in detail below.

#### Rolling MinHash

Using the fact that consecutive windows of the genome are related, we developed a *rolling* MinHash technique to more efficiently compute the fingerprint of window *w*_*p+1*_ from that of window *w*_*p*_, where *p* and *p + 1* are consecutive positions in the genome. This technique is applied as follows.

Let *M* be a matrix of size *L* × *n*, where *L* is the length of the fingerprint and *n* is the size of the kmer set. The function MinHashRoll shown in Algorithm 1 takes as argument the matrix *M*, the column index *m*_*oldest*_, and the fingerprint vector *F*_*w*_ obtained for window *w*_*p*_; it then updates the state of these three variables accordingly, with the fingerprint of *w*_*p+1*_ being stored in *F*_*w*_. The three variables are initialized for the first genome window *w*_*0*_ as follows: *M*(*i,j*) = *h*_*i*_(*w*_*o*_*j*__), where *w*_*0*_*j*__ is the kmer at position *j* in *w*_*0*_, *m*_*oldest*_ = 0, and *F*_*w*_(*i*) = min(*M*(*i*, ·)). The **MinHashRoll** procedure is then applied for all successive windows *p* > 0. The optimization is due to the fact that the minimum over each row of *M* only needs to be recomputed when the *w*_*p*_ minimum value in that row comes from a column corresponding to *m*_*oldest*_ and is smaller than the hash value of the last kmer in *w*_*p+1*_. This technique greatly reduced the computation time of the indexing step.

#### Window Bucketing

By the MinHash property the number of entries shared between the fingerprints of two given sequences is equivalent to the similarity of the original sequence sets, near-duplicate sequences resulting in fingerprints sharing multiple ƒ_i_ entries; therefore, in order to identify such sequences, we need to efficiently compare their corresponding fingerprints. Since pair wise fingerprint comparisons would result in an infeasible running time, the fingerprints are further hashed into *T* hash tables using LSH as follows. Let *proj*_*F*_ be a projection of the fingerprint vector *F* onto 6-dimensions. Given *b* and *T*, we generate *T* random projections of F by creating *T* vectors of length *b* of sparse indices into *F* and extracting the *F* coordinates at those indices. Each projection vector is then hashed into *B* buckets (s.t. *B* = 2^*M*^) using the multiply-shift hash function *Ĥ*_*V*_ initialized with a vector (*a*_*0*_,…, *a*_*b-1*_) of random odd 64-bit integers. The hash value of each projection is computed as follows: 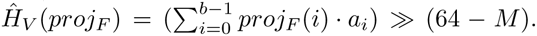. As a result, each of the *B* buckets in a given hash table, will store the positions of the windows whose fingerprints under the table’s projection hashed to that bucket (i.e., windows that share at least the *b* fingerprint coordinates selected by the projection). The index construction is schematically illustrated in Figure 1.

**Fig. 1:**
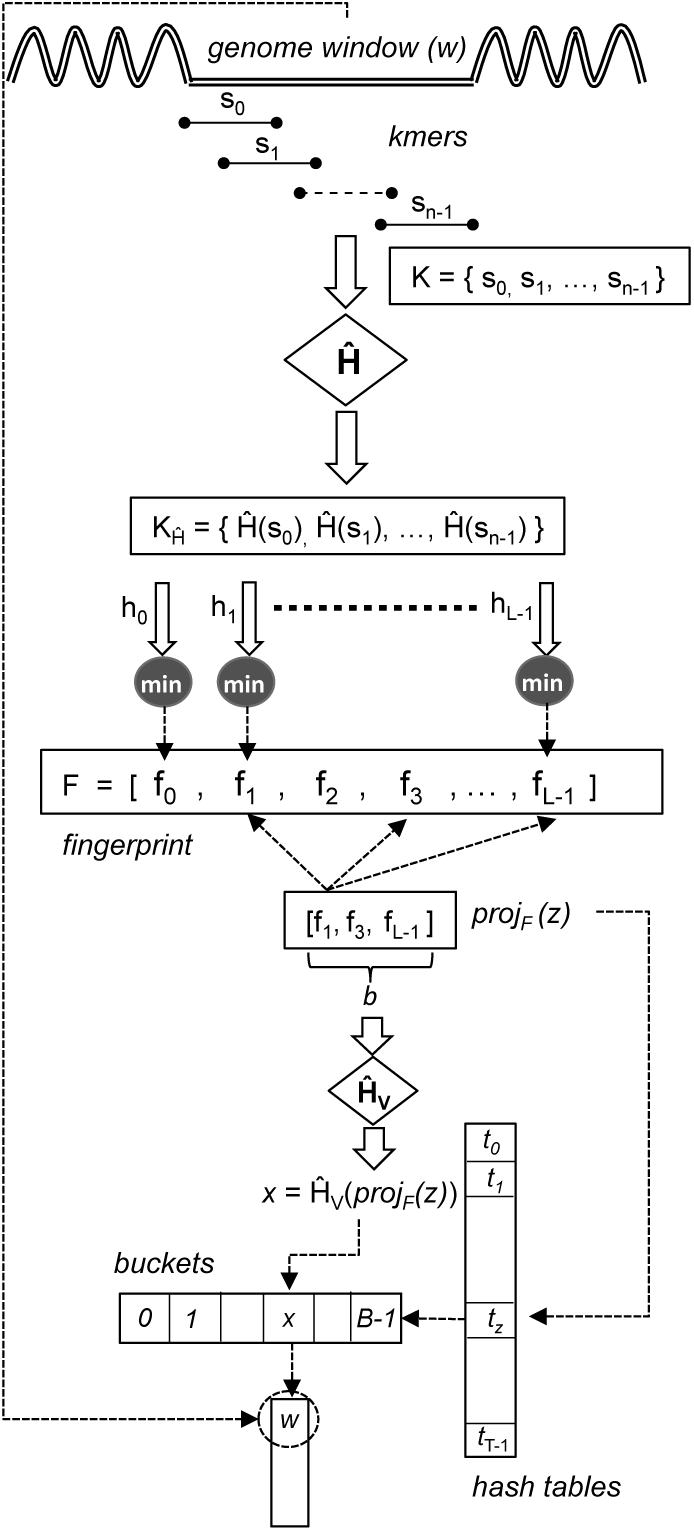
Indexing of a single window of the genome using *L* hash functions for the fingerprint, *T* hash tables, and projections onto *b*=3 dimensions.

#### Memory Reduction and Parallel Construction

There is a clear tradeoff between the number of hash tables used and the sensitivity of the algorithm, more hash tables allowing us to identity more potential hits. However, the space consumption scales linearly with the number of tables and when indexing the full human genome, the memory requirement for large enough *T* can be too high. In order to reduce the memory footprint of the index, we use the following scheme: instead of storing each position separately in the buckets, we store *contigs*, defined by a position and a length, that represent consecutive genome windows that hashed to the same bucket under the same projection. Since the genome is indexed in a linear fashion, the positions will be inserted into the buckets in sorted order; therefore, it is only necessary to check the last inserted position. As a result of this storing scheme, the produced indices are significantly smaller in size. The greatest compression is achieved for larger windows. For example, for window length of 150-bp, 2.83B genome windows were found to have valid MinHash fingerprints; bucketing them into *T* = 78 tables resulted in 6.58B total entries across all the tables, which constitutes a 33.5x size reduction. For windows of length 1000-bp, the reduction in size was 245 x. The memory requirements for the full human genome reference GRCh37 MHG index under the parameters *L* = 128, *T* = 78, *b* = 2, *M* = 18 used in our experiments vary according to the index size as follows: 150-bp: 74GB, 350-bp: 30GB, 1000-bp: 11GB, 10000-bp: 1.1GB. To further reduce the index size, we also provide optional sampling, under which the position *p* is not added to the bucket if it is within *∈* of *p′*, which can result in potentially larger contigs to be handled during alignment (this behavior is disabled by default).

We parallelized the index construction algorithm using OpenMP and use the TBB C++ library’s scalable allocators [28] to optimize the frequent bucket memory allocations. Each thread hashes the windows of a contiguous region of the reference genome sequence and maintains local buckets to eliminate lock contention. When the threads are done processing the genome, the per-thread local buckets are merged. Given the present scheme, indexing the full reference genome on the Intel Xeon X5550 processor takes 1.3 hours using 4 cores on windows of 1000-bp with the parameter setting *L* = 128,*T* = 78,*b* = 2,*M* = 18.

## 3.3 Secure Read Mapping on Hybrid Clouds

### Overview

At a high level, our alignment algorithm can be divided into two main steps: (1) read MinHash fingerprint computation to select candidate contigs from the precomputed MinHash reference index (Phase 1) and (2) kmer voting to select the optimal alignment position among the candidate contigs (Phase 2). Phase 1 constitutes a small fraction of the runtime for typical NGS datasets and currently runs on the secure client, while Phase 2 is outsourced to the public cloud. In order to outsource Phase 2, the client encrypts and transfers kmers generated from the read and candidate windows to the cloud, which can then be securely used for voting. We discuss the algorithm and privacy analysis of each phase below.

### Threat Assumptions

We assume the “honest-but-curious” adversary cloud model, which follows the read alignment protocol correctly but might still try to infer the read sequences from the input data it sees and any background information it has obtained. The computation on the private client and the client-server communications are assumed to be secure. Our goal is to prevent the cloud from deciphering the original read sequences, which can lead to the identification of the individual they came from.

We reasonably assume that the adversary has access to the standard reference genome used during alignment. This data enables the adversary to perform numerous statistical attacks even on encrypted kmer datasets if a deterministic cryptographic scheme (DET) is used to encrypt the kmers, which will produce the same ciphertext value for equal plaintext kmers, allowing the adversary to detect repeats. We differentiate between local repeat structures (LRS), formed by equal kmers inside each read or contig, and global repeat structures (GRS), formed by kmer collisions across reads or contigs. The information leaked by both types of repeat structures can be exploited in a variety of attacks.

For instance, an LRS attack counting intra-read or contig kmers can be designed as follows. For each window of the genome (e.g. of length equal to the read length), the adversary can precompute kmer histograms representing how many times different kmers occurred in this window. Using LRS, similar histograms can also be computed for each read from the encrypted kmers. Finding matches between read and reference histograms can then potentially uncover the encrypted read sequences. We instrumented such an attack in order to assess how identifiable are the reference windows from their kmer histograms. For a window length of 150-bp, we found 91% of the windows of the genome to have no 20-bp long repeat kmers (i.e., every kmer occurs exactly once, making such windows unidentifiable); however, various unique patterns emerged in the remaining 9% of the windows, suggesting that leaking LRS is not secure.

Similarly, the GRS information can also be used to instrument a frequency attack by counting how many times each kmer occurred in the entire read dataset. Assuming that this frequency is equivalent to the kmer frequency inside the reference genome, the adversary can create a mapping between plaintext sequences and encrypted kmers found to occur with similar frequencies (this attack was analyzed by Chen et al [8] since their algorithm allows the adversary to compute such frequencies; however, they determined that this information would not offer much re-identification power to the attacker). GRS information can be further exploited if the relative positions of the kmers inside each sequence are also known. For instance, the adversary can attempt to assemble reads or initial contigs into longer contigs by detecting suffix-prefix kmer overlaps (e.g., using a standard overlap graph assembly algorithm [4]). Longer contigs can then enable more effective histogram attacks since their kmer histograms would result in signatures that are more unique (especially when coupled with kmer position information). Furthermore, identifying repeating kmer tuples in non-overlapping reads or contigs can also leak information due to potential unique co-occurrence patterns (we refer to attacks using this information as GRS-pattern attacks). For example, if two kmers are observed in one contig at a relative distance x but in a different contig at a relative distance y, then if there exists a single pair of kmers that co-occur at distances x and y in the reference genome, we can exactly identify these two kmers. Any number of more complex co-occurrence patterns can be similarly exploited.

Given the above vulnerabilities, our voting technique must not leak **LRS** and GRS information. In order to provide this guarantee, we apply a masking scheme to hide local repeats and employ different encryption keys for each voting task to hide GRS. We provide further details and analyze the security guarantees of our approach in the next sections.

### Phase 1: Read MinHash Fingerprinting and Candidate Contig Identification

During the first phase of the alignment process we compute the MinHash fingerprints of each read in order to identify a small set of candidate alignment positions in the genome, which are expected to be similar to the reads under the Jaccard similarity metric. The fingerprints are computed using the procedure described in Section 3.1. Given the fingerprints, we also calculate the *T* b-dimensional projection hash values to retrieve the genome windows that share the fingerprint entries selected by each projection. As detailed in Section 3.2, each of these hash values corresponds to a particular bucket in the *T* index hash tables. Thus we obtain *T* buckets, each containing contigs of windows which share at least *b* entries with the read (we use b=2 in our experiments, which guarantees that the candidate windows and the read share at least two entries in their fingerprints). This step comprises only a small portion of the total alignment runtime for typical read lengths (see Results) and is not currently outsourced to the public cloud in order to avoid the communication costs associated with shipping the read kmers, as well as potential inference attacks (see Appendix A).

Intuitively, the more buckets a window shares with the read, the more likely it is to be the correct alignment: a perfect match, for instance, would share all *T* buckets. Therefore, given the contigs from all the buckets, the final processing step in identifying the best candidate contigs is to find the rough contig duplicates and count the total number of buckets they are present in, *b*_*hits*_. To do so, our client performs a simple n-way merge using a min-priority heap that stores (contig, *b*_*id*_) tuples of the entries in the buckets (where *b*_*id*_ uniquely identifies each bucket), relying on the fact that the buckets have been sorted by increasing position during indexing. We use a threshold *b*_*min_–_hits*_ to represent the minimum *b*_*hits*_ required for a contig to be considered a strong candidate, as well as keep track of the highest *b*_*hits*_ seen so far, *b*_*best_–_hits*_. A given contig is passed to the next alignment step if its *b*_*hits*_ ≥ *b*_*min_–_hits*_ and it is within some predefined range from *b*_*best_–_hits*_. Figure 2 illustrates the effectiveness of the MinHash index in reducing the number of candidate alignment positions to only a few contigs per read given the *b*_*min_–_hits*_ thresholds of 1 (i.e. the contig is processed even if it is only present in one bucket) and 2 (i.e. the contig must be present in at least two separate buckets). As expected, a higher *b*_*min_–_hits*_ threshold results in a much smaller number of processed contigs. We can see that our MinHash index allows us to select fewer than 10 contigs for most reads. This high selectivity is crucial for our alignment algorithm, since it significantly decreases the overheads associated with encrypting and transferring the selected contigs for Phase 2.

**Fig. 2:**
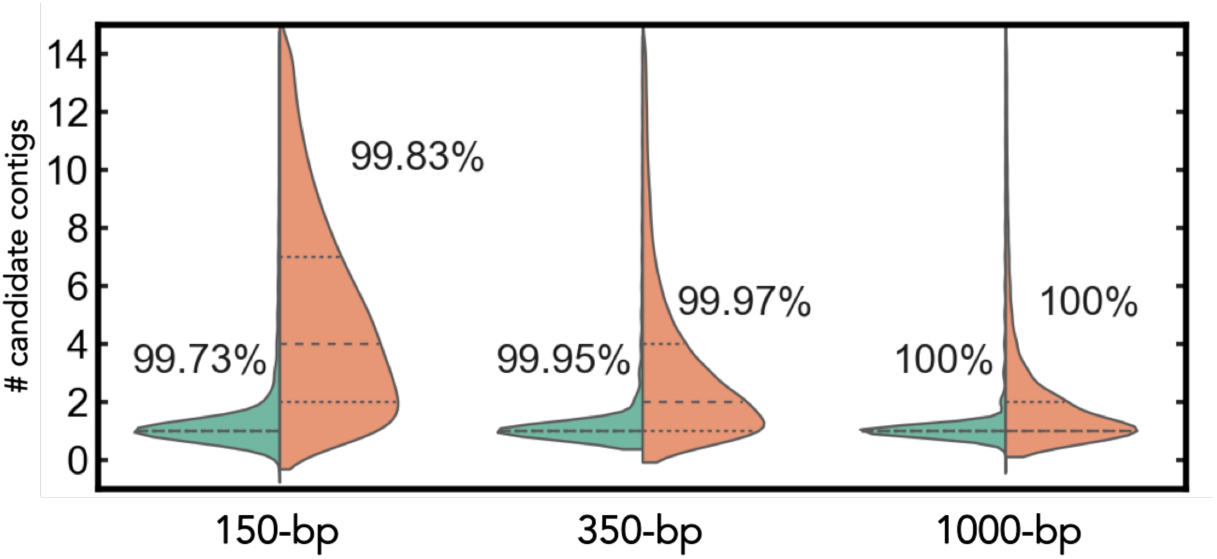
Distribution of the number of candidate contigs (cropped) selected per read computed over 100K read datasets for varying read lengths. The percentage of reads for which the true contig was selected is shown on top of each distribution.

### Phase 2: Secure Kmer Voting

Given a contig, the second phase of the algorithm is to estimate the displacement ϕ which best aligns it to the read, as well as a *score* which measures the quality of that alignment. We define a *voting task* to represent such a read and contig pair and employ the following voting technique to process each task resulting from Phase 1 (each task can be executed in parallel on the cloud).

Let’s start by representing a read *R* and a candidate contig *C* as a set of kmers and their associated positions: 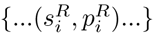 and 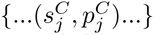 respectively (our implementation currently uses 20-bp overlapping kmers). Assume we have found the correct contig. With no error and only unique kmers, a single match 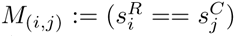 would unambiguously yield the correct alignment. Namely: 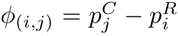. We say that match *M*_*(i,j)*_ *votes* for the alignment position *ϕ*(_*i,j*_). Given noisy data and non-unique kmers, however, no single match is guaranteed to be correct. Furthermore, due to potential insertion and deletion errors, all correct matches may still not agree on a precise alignment. Therefore, we search for the alignment which (roughly) explains the maximum number of kmer matches as follows. First we let each kmer in the read matching a kmer in the contig vote for the respective alignment positions. As a result we obtain a vector *V* of votes for each position in the contig. Since we expect some votes to be split across neighboring positions due to indels, we perform a simple convolution over *V* to collect neighboring votes, resulting in a new vector 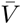, where 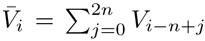 (we use *n* = 10 in our experiments). The position in 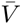 with the maximum number of votes is selected as the best alignment position, with the number of votes representing its score (if a range of positions have the maximum number of votes, we pick the middle position in the range). As we process all the contigs, we keep the best score *r*_*1*_ and second best score *r*_*2*_ found per read in order to identify situations when a read can map well to multiple positions and estimate the mapping quality of our alignments. We currently use the simple formula 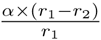 (similar to other methods like BWA-SW [22]) for our mapping quality.

Our goal is to outsource Phase 2 securely to the cloud. The main requirement then is to encrypt the kmers and positions securely. We propose two protocols for executing the voting scheme above, using DET and HOM to encrypt the kmers, respectively. Our implementation uses the first protocol since it is more practical and we focus on this protocol in the rest of the manuscript. The second protocol is described in Appendix B.

### Kmer Hashing

A key property of our voting protocol is that it does not require the comparison of kmers across voting tasks, since we only need to detect kmer matches between a read and its candidate contig to count the votes. This allows us to avoid leaking GRS information by associating unique secret keys with each voting task and computing the kmer hashes as follows. First we apply a standard cryptographic hash function SHA-1 (with a 256-bit secret key *K*) to each kmer, which guarantees that even kmers that only differ by a single nucleotide are hashed to completely different values. SHA-1 produces a 20-byte digest, of which we use the first 8 bytes as our kmer hash (note, both the SHA-1 hashing scheme and the length of the kmer hash can be easily replaced in our method by another deterministic cryptographic hashing scheme without affecting its resulting alignment accuracy). Next we apply an additional encryption step to each digest using a set of two randomly-generated 64-bit keys, 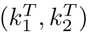, that are unique for each voting task T, such that a kmer s_i_ associated with a given task is set to the following value: 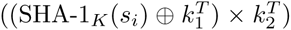 While we could use a unique per task secret key with SHA-1 to avoid the second encryption step, this would prevent us from precomputing the contig kmer hashes ahead of time, increasing our voting data encryption overheads on the client. Instead, using the two-step procedure, we can optimize the encryption time by only having to apply the faster second step to the precomputed SHA-1 reference kmers. As a result, kmer collisions can be detected within each task but not across different tasks. Finally, in order to avoid leaking LRS information, we need to mask repeat kmers inside each read and contig. To mask a repeat kmer, we can replace its hash value with a random 64-bit value. While this would be sufficient when comparing kmer values only, such masking can be detected when evaluating the matching pattern between read and contig kmers. For example, if kmer positions are given and we observe a kmer mismatch enclosed by kmer matches in both the read and the contig, we can deduce that the mismatch is due to masking (since the matching predecessor and successor kmers will entirely cover the bases of the mismatching kmer). Therefore, in order to avoid this leak, we mask the neighboring kmers of each repeat as well, such that there are at least *v* masked kmers around each repeat, where *v* is the length of the voting kmers.

In order to minimize bandwidth, our method also supports *sampling* the contig kmers, such that only a fraction of kmers are sent to the server and participate in voting. The configurable sampling rate parameter, *ρ*, controls the sparsity of the sampled kmers. We evaluate the effects of sampling and repeat masking on alignment accuracy in the Results section.

### Position Binning

While we cannot reveal the actual position of contig kmers in the reference genome, we only need the relative positions of the kmers inside the contig in order to compute the local displacement of each match and count the votes. The displacement that received the most votes could then be sent to the client and added to the global start position of the contig inside the reference. Our protocol reveals only the partial ordering of the kmers inside the read and contig, shuffling kmers falling into different bins. In our experiments, we use a bin size *β* = 20 (equal to the length of the kmers), such that the position of each kmer is ambiguous within *β* positions and only the relative positions of the bins are known (e.g. *β* = 1 would reveal the full ordering of the kmers). More specifically, the stream of read or contig kmers received at the server, can be broken down into bins of size *β*, such that bin Bj (i.e. *i*th bin in the stream) contains *β* kmers in an arbitrary random order that fall in the range [*l, h*) = [*i* × *β*, *i* × *β* + *β*). In this case, each kmer match casts a vote for a range of positions (as opposed to a single position described above) equivalent to the positions obtained if every single kmer in the contig bin matched every kmer in the corresponding read bin, namely: [*l*^*C*^ – *h*^*R*^ + 1, *h*^*C*^ – *l*_*R*_].

When kmer sampling is enabled (as described above), binning the positions can mediate the effect of masking repeat neighborhoods on alignment accuracy (as shown in Results). Namely, for a sampling rate *ρ*, if there are *β/ρ* unique kmers in the bin, these kmers will be selected for voting (without having to reveal their precise positions in the bin) and the bin will not be masked; otherwise, masking will be applied on the entire bin. Furthermore, binning the positions minimizes the information learned about the pattern in which the read kmers matched against the contig.

### Privacy Analysis

As discussed above, our voting scheme does not leak LRS and GRS information, preventing the adversary from detecting any repeats in the dataset and thus avoiding the defined frequency attacks. The main information the adversary has is the result of each voting task: the number of kmers matched and their binned positions. However, given the presence of errors across the reads (a single error will result in a different matching pattern) and SNPs in the genome (e.g., unrelated reads covering different SNPs might share the same voting results), we are unaware of a statistical attack that could use this information. Additionally we also reveal the length of the candidate contigs. The length of the contig is determined by how many contiguous windows of the genome fell into the MHG buckets matched by a read’s MinHash fingerprint. While precomputing the same bucket structure would be difficult for the adversary when the projections and the *L* hash functions are unknown, we can eliminate this vulnerability by splitting long contigs into multiple pieces up to length *∧*. Furthermore, we also add a small randomly selected amount of padding to each contig in order to prevent contig matching across tasks using their lengths. Splitting the contigs or adding extra padding does not affect the accuracy of the results, while having a negligible compute and communication overhead (since the amount of padding used and the number of required splits is small).

By default, our voting tasks consist of a single read and candidate contig pair. This prevents the adversary from learning the number of candidate contigs matched by each read (since it cannot correlate which tasks belong to the same read), as well as protects against the GRS-pattern attack. While most reads match only a few contigs (see Figure 2), this number can vary based on how repetitive the read sequences are in the genome (and can represent a strong signal for highly repetitive sequences); however, below a certain low threshold *Z*, the number of reads matching at least *Z* candidate contigs would become uninformative. Therefore, we provide functionality for *batching* contigs, such that a task can store a read and multiple candidate contigs. This mode can reduce the encryption time and the bandwidth requirement (since we would not need to ship read kmers encrypted with different task keys for every single candidate contig). However, batching does result in vulnerability to the **GRS-pattern** attack. In order to evaluate our method with maximum security guarantees, we do not enable batching in our experiments.

## 4 Results

First it is important to assess whether our approach of combining LSH and kmer voting is competitive in accuracy and performance to the existing read aligners. In order to do that we simulated 150-bp and 350-bp long read datasets of 100K reads each from the human genome reference GRCh37 using the wgsim program [19] with the following (default) parameters: SNP mutation rate = 0.09%, indel mutation rate = 0.01%, and sequencing base error rate = 1% and 2% (typical for the Illumina sequencing datasets). We compare our results to two of the most popular and efficient read aligners, BWA-MEM [20] and Bowtie2 [18] using their default settings. All the experiments were performed on a single 2.67 GHz Intel Xeon X5550 processor. The accuracy results in a distributed setting will remain the same, while some additional costs will be incurred due to the client-cloud communication. Runtime is reported for single-threaded execution. We report the percentage of reads aligned with a mapping quality greater than 10 (Q10) and the percentage of incorrect alignments out of the Q10 mappings (Err). We considered a read to be mapped correctly if its reported position was within 20-bp of the true position.

The results in Table 1 and Figure 3 demonstrate that **BALAUR** performs well in comparison to other aligners, achieving similar high percentages of Q10 mapped reads and low error rates (with BWA-MEM having a lead over the other aligners). Examining the Q10 incorrect mappings, we found that many of them typically occur when the true position of the read has not been evaluated in the second phase of the alignment process, while another alternative position was found to match the read fairly well (e.g. in the case of a repeat). Setting *b*_*min_–_hits*_ to 1 and disabling the filtering based on b_best hits_, which would examine every single contig that fell into the read buckets, or increasing the index size, which would increase the likelihood of the true position being in any of the read buckets, can be expected to increase accuracy; however, at the expense of slowing down the runtime. The **BALAUR** results were obtained with MHG index parameters: L=128, T=78, b=2, M=18. We also used a kmer sampling rate of *ρ* = 3, position bin size *β* = 20, and no contig batching (i.e. voting tasks composed of a read and a single contig only). We do not incorporate initialization costs (e.g. the allocation of kmer transfer buffers and MHG loading) into the reported runtime. To examine the effects of LRS kmer masking and position binning, Figure 3 also shows the results for **BALAUR-vanilla**, which is a non-secure version of our algorithm using a faster non-cryptographic hash function (namely, CityHash64 [1]), no repeat masking, and no position binning. It can be seen that for the above datasets, kmer masking and position binning does not have a significant impact on the accuracy of the alignment results. Further evaluation of these settings and the effect of kmer sampling is shown in Supplementary Figure 1.

**Table 1:**
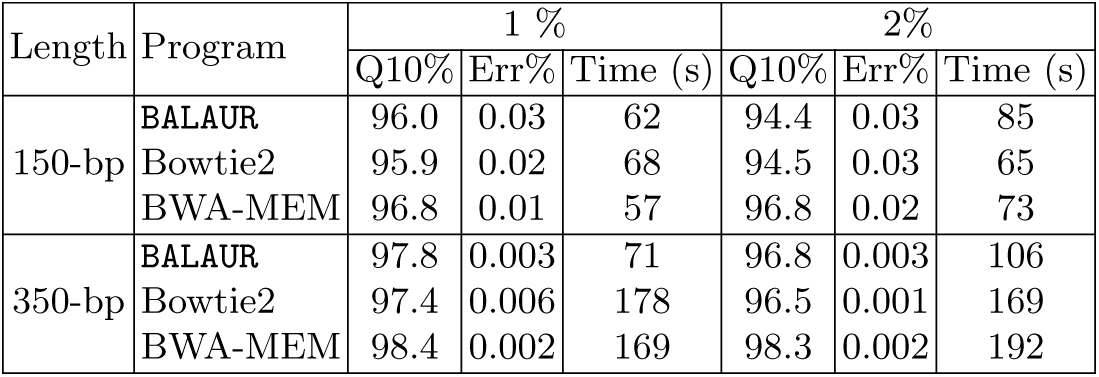
Evaluation on simulated whole-genome GRCh37 reads.

**Fig. 3:**
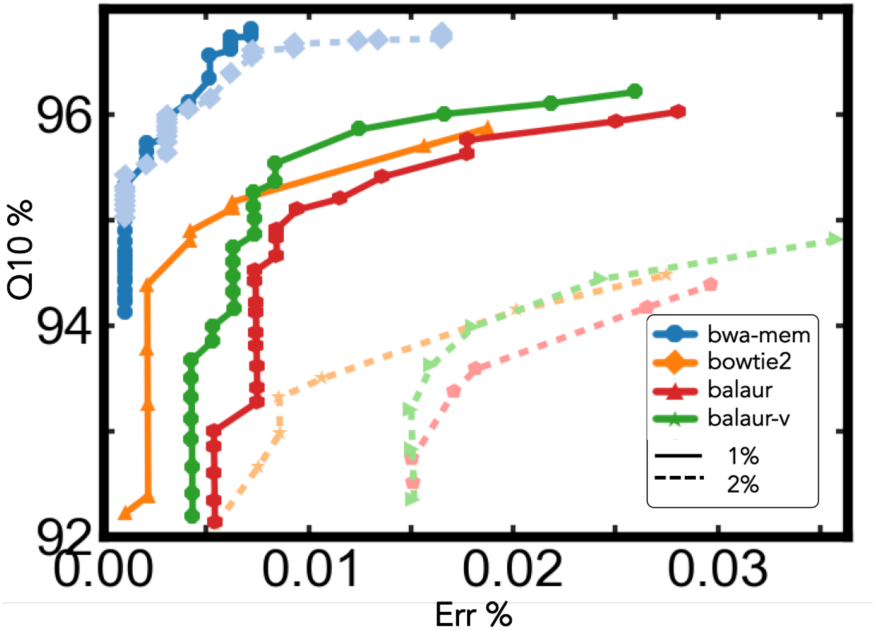
Evaluation on 150-bp simulated reads.

### Long Reads

As suggested by the results on the 350-bp long read dataset, **BALAUR** can be substantially faster than other aligners on longer read lengths. Therefore, orthogonally to security guarantees we also explore its effectiveness as a long read aligner. Several sequencing technologies can currently produce low error long reads; for example, the PacBio CCS reads are ≈3-kb long reads generated by the single molecule real-time (SMRT) PacBio RS platform with an approximate 2.5% error rate [16]; similarly, Illumina TruSeq synthetic long reads have an estimated error rate of only 0.0286% per base [26]. However, typical long read datasets contain higher error rates. In order to evaluate how our aligner scales with increasing error rates, we simulated 1,000-bp and 10,000-bp read datasets of 100K reads from the human genome reference GRCh37 using wgsim [19]. We used the wgsim default polymorphism parameters and set the base error rate to 1% and 2% for the 1,000-bp dataset and 4%, 8%, and 10% for the 10,000-bp reads, respectively. Figure 4 and Supplementary Table 1 present the results. We compare the results on the 10,000-bp reads with the BWA-MEM and BWA-SW [22] aligners only here since Bowtie2 took a significantly longer time to run on this data. For this experiment we used **BALAUR-vanilla**, kmer length of 32-bp, and varying sampling rates and MHG index sizes (the specific configurations are presented in Supplementary Table 1). It can be seen that **BALAUR-vanilla** achieves significant speedups (ranging from 4-40×) over the two aligners, while maintaining high accuracy.

**Fig. 4:**
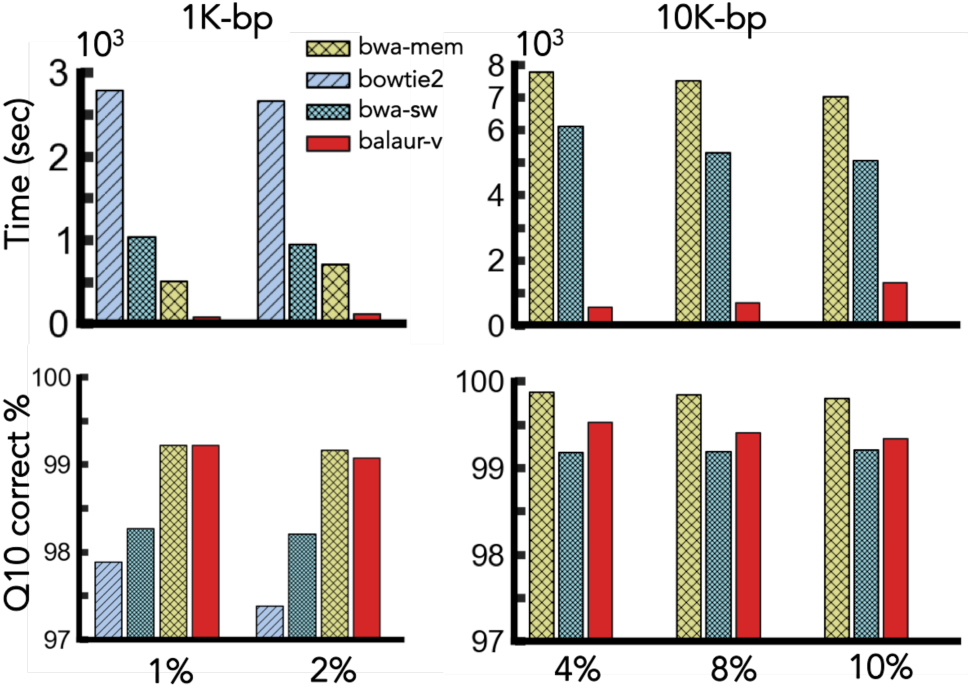
Evaluation on simulated long reads with varying sequencing error rates of 1 - 10%.

### Real Reads

We also assessed the performance of **BALAUR** on the following two real read datasets: (1) 1M 150-bp HiSeq2500 reads of the NA12878/HG001 genome and (2) 1M 150-bp HiSeq2500 reads of the NA24385/HG002 genome (with the same parameters used for simulated read experiments). The results are shown in Figure 5. Since we don’t know the true alignment position of each read, we report the percentage of Q10 mappings and the runtime. As in simulations, **BALAUR** achieves similar Q10% results; furthermore, we have found that more than 99.95% of its Q10 mappings are within 100-bp of the BWA-MEM mapping results.

**Fig. 5:**
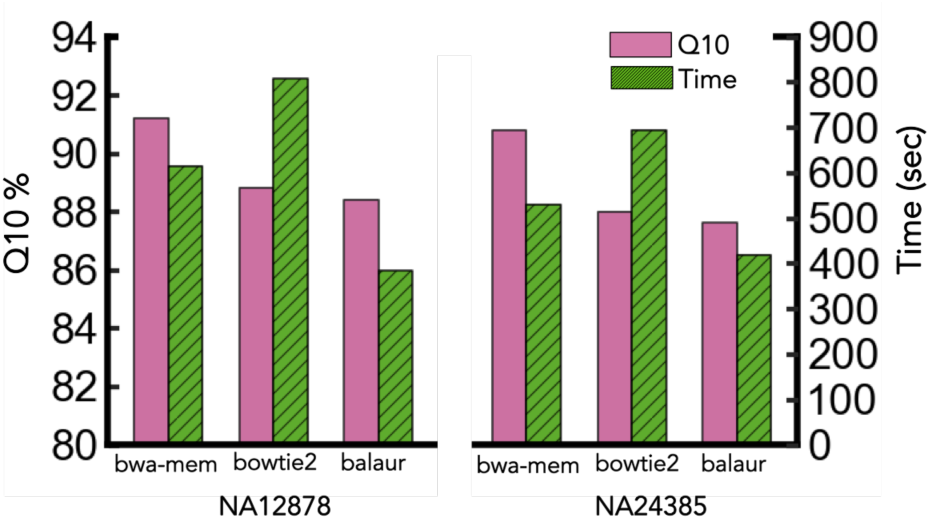
Evaluation on real read datasets.

### Runtime and Communication Cost Breakdown

Finally we evaluate how much computation can be outsourced to the cloud for each read dataset and the costs associated with each alignment phase. Figure 6 and Supplementary Table 2 present the breakdown of Phase 1 and Phase 2 costs and the required bandwidth for each examined dataset. The reported “encryption” time includes both kmer hashing and repeat masking, while the “reporting” time represents the post-processing of the voting results on the client (i.e. selecting the best alignment positions and SAM output file I/O). It can be seen that the voting step, which can be securely outsourced to the cloud, accounts for a high percentage of the total runtime, varying across read lengths and sequencing error rates. Since by design our algorithm must send both the encrypted read and contig kmers as part of each voting task, we must ensure that our protocol imposes practical communication overheads. From this consideration, the LSH step is critical since it allows us to narrow down the number of contigs (and hence kmers) sent per read. In the simulated 150-bp 1% read length experiment, we sent 2.8M contigs total for the read dataset; together with the read kmers (with 64bits of data per kmer) this required 4.9GB bandwidth. We find this cost to be high but still practical for current networking speeds, although additional heuristics need to be explored to drive down the cost further; for example, it would take 40 seconds to transfer this data on a fast 1 Gbps link. Moreover, this communication delay can be hidden by overlapping the data transfer with computation at the client side (i.e. the client can encrypt the next batch of reads while a previous batch is being transmitted).

**Fig. 6:**
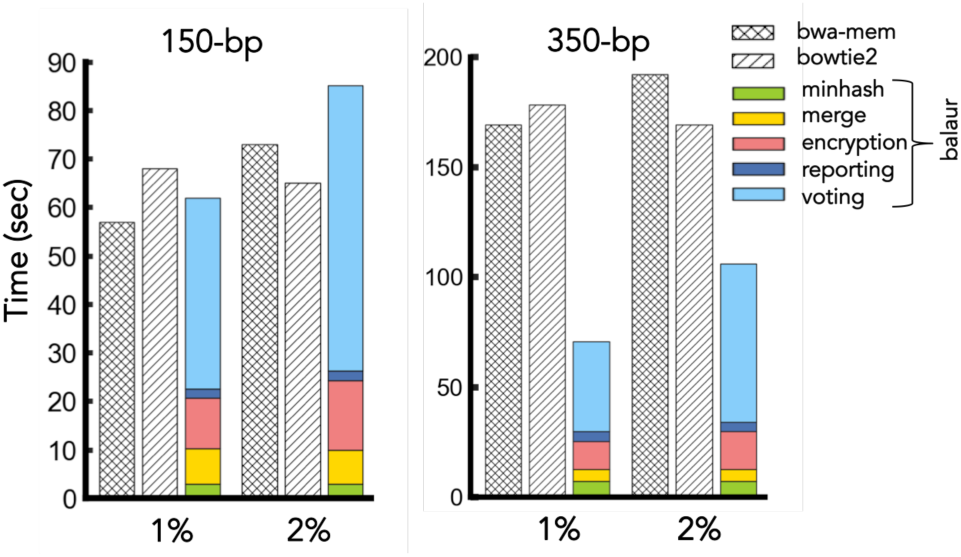
Runtime breakdown.

## 5 Conclusion

**BALAUR** is a practical secure read-mapping algorithm for hybrid clouds, based on LSH and kmer voting, and is highly competitive with non-cryptographic state-of-the-art read aligners in both accuracy and performance. It incorporates an efficient technique to index the reference genome using the MinHash algorithm, which is effective in reducing the computation and communication costs of its secure voting scheme. Orthogonally, it is highly effective for long read alignment, achieving significant runtime speedups over existing tools on longer read lengths. In this paper we analyzed the security guarantees of our approach, describing what information it leaks to the cloud and addressing any adversarial attacks known to us. We leave several interesting directions for future work. In particular, currently our MinHash and voting schemes use a simple kmer generation principle (i.e., all overlapping kmers of a certain length); however, more sophisticated kmer designs could be explored to improve sensitivity (e.g., sparse patterns). Furthermore, a statistical analysis of the risks associated with batching multiple contigs per voting task can be performed to devise an optimal batching scheme, which minimizes the number of tasks per read (thus decreasing the bandwidth requirements) while maintaining appropriate security guarantees.

## Acknowledgement

We thank Stephen Miller and Valeria Nikolaenko for their useful feedback and advice.

## Appendix

**Supplementary Table 1:**
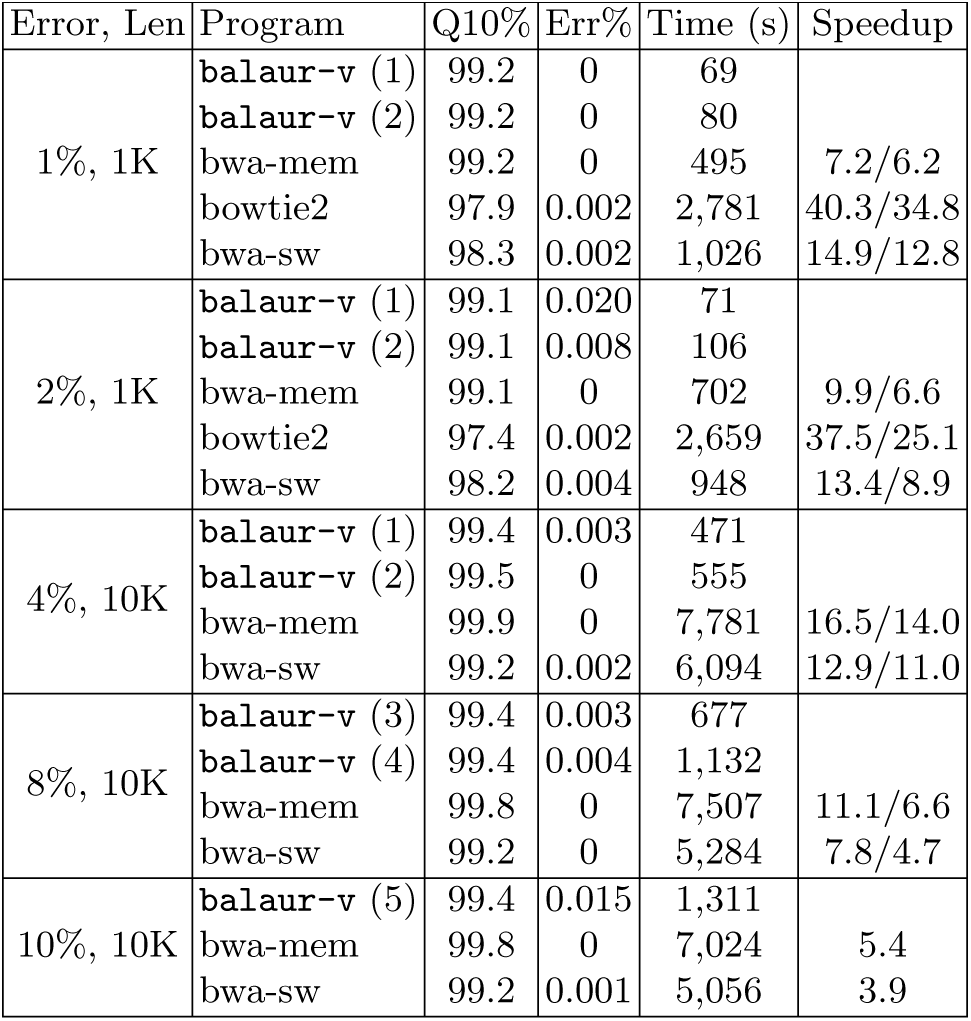
Evaluation on simulated long reads. (1) MHG0, *b*_*m*_ = 2; (2) MHG0, *b*_*m*_ = 1; (3) MHG2, *b*_*m*_ = 4; (4) MHG1, *b*_*m*_ = 1; (5) MHG2, *b*_*m*_ = 1. *ρ* = 4 (1K); *ρ* = 8 (10K, 4%); *ρ* = 2 (10K, 8% and 10 %). *b*_*m*_ stands for *b*_*min-hits*_. MHG0: (L=128, T=78, b=2, M=18); MHG1: (L=256, T=512, b=2, M=16), 6.7GB; MHG2: (L=256, T=1024, b=2, M=16), 14GB.

| Error, Len | Program | Q10% | Err% | Time (s) | Speedup |
| --- | --- | --- | --- | --- | --- |
| 1%, 1K | **balaur-v** (1) | 99.2 | 0 | 69 |  |
|  | **balaur-v** (2) | 99.2 | 0 | 80 |  |
|  | bwa-mem | 99.2 | 0 | 495 | 7.2/6.2 |
|  | bowtie2 | 97.9 | 0.002 | 2,781 | 40.3/34.8 |
|  | bwa-sw | 98.3 | 0.002 | 1,026 | 14.9/12.8 |
| 2%, 1K | **balaur-v** (1) | 99.1 | 0.020 | 71 |  |
|  | **balaur-v** (2) | 99.1 | 0.008 | 106 |  |
|  | bwa-mem | 99.1 | 0 | 702 | 9.9/6.6 |
|  | bowtie2 | 97.4 | 0.002 | 2,659 | 37.5/25.1 |
|  | bwa-sw | 98.2 | 0.004 | 948 | 13.4/8.9 |
| 4%, 10K | **balaur-v** (1) | 99.4 | 0.003 | 471 |  |
|  | **balaur-v** (2) | 99.5 | 0 | 555 |  |
|  | bwa-mem | 99.9 | 0 | 7,781 | 16.5/14.0 |
|  | bwa-sw | 99.2 | 0.002 | 6,094 | 12.9/11.0 |
| 8%, 10K | **balaur-v** (3) | 99.4 | 0.003 | 677 |  |
|  | **balaur-v** (4) | 99.4 | 0.004 | 1,132 |  |
|  | bwa-mem | 99.8 | 0 | 7,507 | 11.1/6.6 |
|  | bwa-sw | 99.2 | 0 | 5,284 | 7.8/4.7 |
| 10%, 10K | **balaur-v** (5) | 99.4 | 0.015 | 1,311 |  |
|  | bwa-mem | 99.8 | 0 | 7,024 | 5.4 |
|  | bwa-sw | 99.2 | 0.001 | 5,056 | 3.9 |

**Supplementary Table 2:** Runtime and bandwidth breakdown by alignment phase.

| Length | Phase 1 Runtime (s) |  | Phase 2 Runtime (s) |  |  | Total Time (s) | Bandwidth |  |
| --- | --- | --- | --- | --- | --- | --- | --- | --- |
|  | MinHash | Merge | Encryption | Voting | Reporting |  | Up (MB) | Down (MB) |
| 150-bp, 1% | 2.9 | 7.4 | 10.3 | **39.3** | 2.0 | 61.9 | 4,930 | 110 |
| 150-bp, 2% | 2.9 | 7.0 | 14.3 | **59.0** | 2.0 | 85.2 | 7,507 | 168 |
| 350-bp, 1% | 7.0 | 5.6 | 12.7 | **40.9** | 4.4 | 70.6 | 4,651 | 44 |
| 350-bp, 2% | 7.0 | 5.4 | 17.2 | **71.8** | 4.4 | 105.8 | 8,323 | 80 |
| NA12878 | 27.7 | 76.4 | 64.3 | **194.4** | 20.3 | 383.1 | 24,613 | 554 |
| NA24385 | 28.7 | 75.0 | 70.6 | **223.0** | 20.5 | 418.0 | 28,423 | 641 |

**Supplementary Figure 1:**
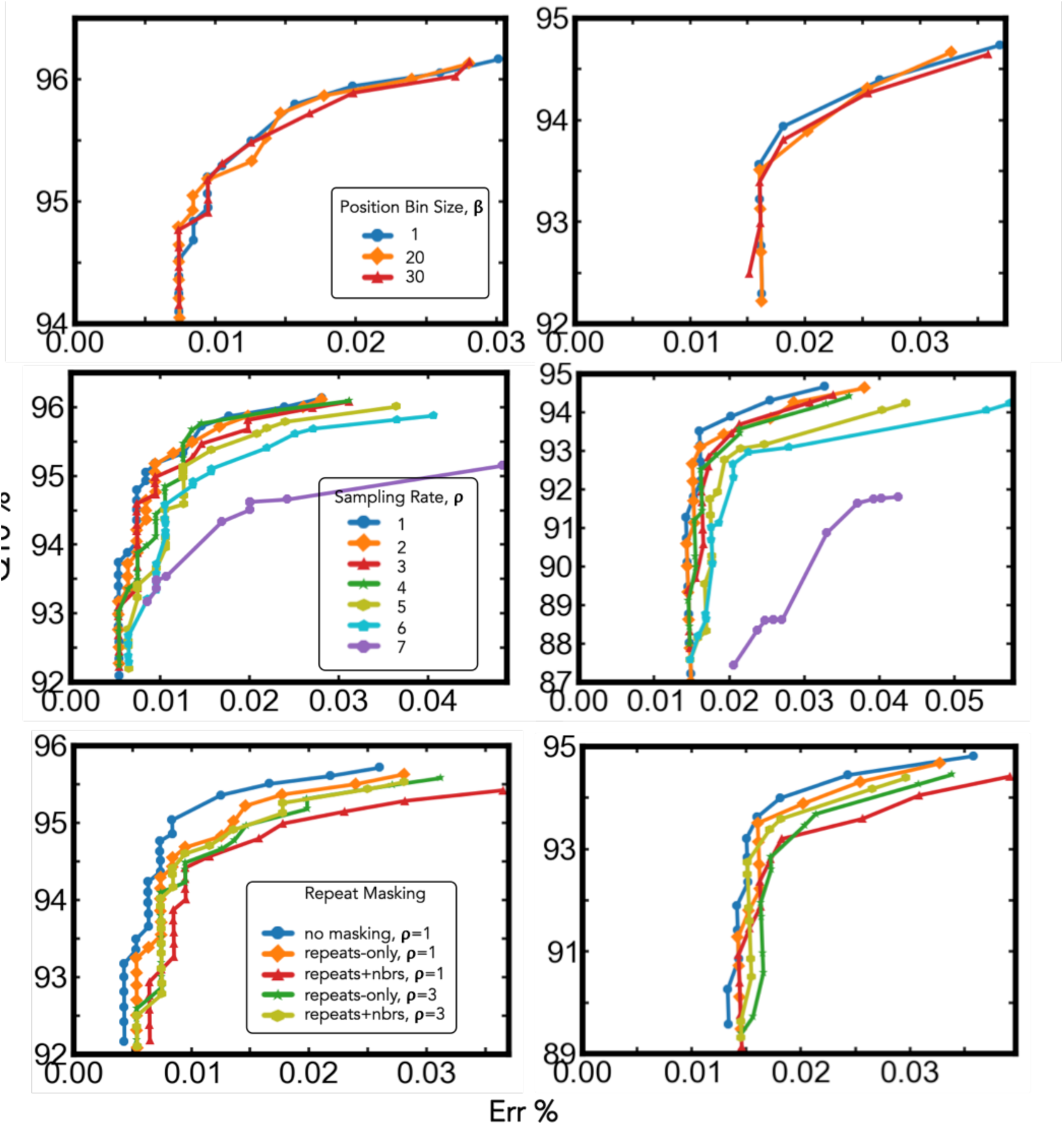
Evaluation of position binning, kmer sampling, and kmer masking on 150-bp 1% (left) and 2% (right) error read datasets.

**A. Privacy Considerations of Phase1: MinHash Fingerprinting**

Although the MinHash fingerprints can be computed from a set of encrypted unique kmers (since the minimum value does not change due to the presence of identical elements in the set or their order), equal kmers across all the reads and the reference must be hashed to the same value to determine set similarities and use the MinHash index. As previously discussed, this would allow the adversary to perform the GRS frequency attack by counting repeats. Furthermore, we would also have to leak kmer linkage information, since the minimum value must be computed with respect to each read kmer set, significantly strengthening the frequency attack. Therefore, our scheme currently avoids exposing this data to the cloud at the expense of performing this step on the client (and of course when reads are very long or *L* is very large, the overhead of this computation will become more significant). There are a couple of possible solutions to remedy the frequency attack here although these are left for future work. The main observation is that the set of features chosen to represent each read and reference window when computing the MinHash fingerprints does not need to be the set of all contiguous overlapping kmers. We can represent a read as a set of more complex features (e.g. sparse kmers, where the sparsity is nondeterministic and unknown to the cloud), making it more difficult for the cloud to compare the frequency of such features with what it can precompute from the reference genome.

**B. Voting Protocol using HOM Kmers and Positions.**

To hide how many read kmers matched the candidate contig, we can encrypt the kmers using a non-deterministic scheme, which produces different ciphertexts for the same plaintexts. In particular, we can use an additive homomorphic encryption scheme (AddHOM), such as the Paillier [27] cryptosystem, to encrypt the kmers and their positions, while still being able to determine whether two kmers matched and what position they voted for upon decryption. AddHOM allows us to find the sum of two plaintext values by computing the product of their ciphertexts. Let 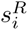 and 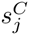 be two plaintext read and contig kmer values, then 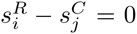 iff 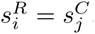. Therefore, we know we have a kmer match if 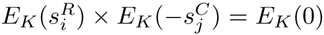 where *E*_*K*_(·) is an AddHOM scheme. Similarly, let 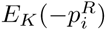 and 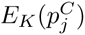 be the encrypted positions of two kmers in the read and contig, respectively, then the encrypted alignment position for which these kmers vote is 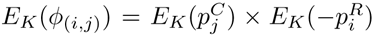. This allows us to formulate the voting protocol as follows. First the client encrypts all kmers and their positions using AddHom and sends them to the cloud. The cloud then computes the tuple 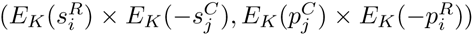 for each pair of encrypted read and contig kmers and sends these back to the client. Finally, the client decrypts each tuple and records its vote when 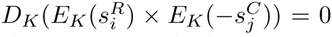, where *D*_*K*_(·) is the decryption procedure. It can then compute *V* and find the position with the maximum number of votes as described above.

While this scheme does not leak any mapping information to the cloud, it increases the complexity of the voting scheme and requires the client to perform elaborate post-processing (including the decryption of the voting results). More specifically, under the DET protocol we can find the kmer matches in O(nlogn) time by sorting the read and contig kmer lists and then merging the results to find the hits. Under the HOM protocol, however, we have to compute and transfer *O*(*n*^2^) pairwise tuples since we can no longer determine the matches on the cloud. This can be mitigated in a hybrid approach that encrypts kmers with DET and only the positions with AddHOM. That being said, the biggest limitation of a scheme using HOM is the additional communication cost it would impose. For example, the Paillier cryptosystem requires 2048 bits per ciphertext, which would result in a highly prohibitive data transfer overhead.

